# Improving night-time sleep with hypnotic suggestions

**DOI:** 10.1101/277566

**Authors:** Maren Jasmin Cordi, Laurent Rossier, Björn Rasch

## Abstract

Slow-wave sleep (SWS) is fundamental for maintaining our health and well-being, and SWS is typically reduced in stress-related sleep disturbances and age-related sleep disorders. We have previously reported that exposure to hypnotic suggestions before sleep effectively increases the duration of SWS during a midday nap in younger and older women suggestible for hypnosis.

However, it remains unclear whether this beneficial effect of hypnosis on SWS can be generalized to night-time sleep and men. Therefore, we tested the effect of the hypnotic suggestions on SWS across an 8 hours night-time sleeping interval in 43 healthy young French-speaking subjects (19 males) of high and low suggestibility. In accordance with our previous results, listening to hypnotic suggestions before sleep increased the amount of SWS in highly suggestible subjects significantly by 13 min compared to a control condition in both genders. Particularly in the first hour, slow-wave activity was significantly increased after hypnosis as compared to the control night in high suggestible. The hypnosis-induced benefits on objective sleep parameters were also reflected in increased subjective sleep quality ratings. Our results demonstrate that hypnotic suggestions are an effective tool to deepen sleep and improve sleep quality also across a whole night of sleep in young healthy men and women. Our findings provide an important basis for the examination and potential application of hypnosis to improve deep sleep in populations with sleep disturbances.

## Introduction

Sleep is critically involved in good health and cognitive functioning [1-3]. However, sleep problems are highly frequent [e.g., 4], which increases the chances of disorders, accidents and deficits in cognitive functioning. For example, Hinz et al. [5] have recently assessed subjective sleep quality in 9284 adults aged 18 to 80 and reported that 36% of the general population indicated having sleep problems. Those troubles were related to fatigue, anxiety, and somatic complaints. For high quality and restorative sleep, the depth of sleep is of particular importance. Deep sleep is called slow-wave sleep (SWS) due to its phenomenological appearance of slow waves in the EEG signal and has been related to cognitive functioning [6], the immune system [7], and mental health [8,9]. Additionally, SWS also is a critical component in the subjective rating of sleep quality [10,11]. However, the prevalence of this sleep stage decreases with increasing age [12] and is often affected by sleep disturbances [13]. SWS is hence a vital component for healthy sleep, and the development and testing of effective and safe methods to protect and improve SWS are highly warranted. We have previously demonstrated that hypnotic suggestions represent an effective means to increase the amount of SWS in a risk-free way [14,15]. Hypnosis has been defined as “a state of consciousness involving focused attention and reduced peripheral awareness characterized by an enhanced capacity for response to suggestion” [16]. The positive effects of hypnosis or suggestions given during a hypnotic state have been demonstrated for several somatic illnesses and as a treatment for smoking cessation and other psychiatric problems [17,18]. Concerning sleep, most studies had reported positive effects on subjective ratings of sleep quality, but were lacking objective evidence for these findings [19,20]. In our two studies we measured sleep with polysomnography during a midday nap and demonstrated that hypnosis increases the amount of SWS from 14 to 23 min in young and from 12 to 21 min in older high suggestible women, respectively. The increases in the amount of SWS were also reflected in an increase in slow-wave activity (SWA) during NREM sleep. Those were the first studies to objectively confirm hypnosis-related changes in sleep architecture. However, both studies were conducted during midday naps in non-habitual nappers, for which homeostatic and circadian sleep pressure should be expected to be low. According to the well-established model of sleep regulation [21], the amount of SWS and SWA strongly depends on both a homeostatic process and a circadian process, resulting in high amounts of SWS when sleep pressure is high due to an extended wakefulness and optimal circadian timing [22]. Thus, it remains unknown whether hypnotic suggestions are also capable of extending SWS and increasing SWA under high sleep pressure conditions, i.e., during normal night-time sleep.

Here we experimentally tested the benefits of hypnotic suggestion given before night-time sleep in healthy young high and low suggestible subjects. Due to the higher sleep pressure, we expected that the effects of hypnosis on SWS would be significant, but lower than in our previous nap studies. In addition, we were interested whether hypnotic suggestion would influence SWS only in the first hour after sleep onset (comparable to the sleep time in our previous nap studies) or also during later periods of sleep. Furthermore, we translated the hypnotic suggestion from German to French and included both female and male participants with a French mother tongue. Thus, we aimed at generalizing our previous findings also from females to males and to another language.

## Materials and Method

### Subjects

We tested a total of 52 healthy, French-speaking younger adults (25 males, between 19 and 31 years, mean age of 22.02 ± standard deviation [SD] of 2.49). Due to technical problems, we could not use sleep data of 5 subjects. Another 4 subjects were excluded due to an amount of wakefulness in at least one of the sessions which exceeded the mean by more than 2 SD. The final sample consisted of 43 participants (19 males, aged 19 to 31 with a mean age of 22.10 ± 2.72 [SD]). The participants refrained from drinking alcohol and caffeine and got up before 8 a.m. on the experimental days. For participation in all sessions, 150 CHF were paid. The Ethics Committee of Lausanne (CER-VD 115-15) had approved the study and all subjects signed the informed consent before participation.

### Procedure

The procedure followed the protocol of the previous two hypnosis studies we performed across a midday nap [i.e., 14,15]. The participants were invited to a total of four sessions, out of which three took place in the sleep laboratory. The first group session included testing subjects’ suggestibility with the standardized questionnaire of the Harvard Group Scale of Hypnotic Suggestibility (HGSHS, German version by Bongartz [23], translated into French by Laurent Rossier). We explicitly informed the subjects about the hypothesis of our study. The second session served as adaptation session in which the participants spent a night (∼10 p.m. to 7 a.m.) in our sleep laboratory attached to the polysomnographic electrodes (i.e., electroencephalography (EEG), electromyography (EMG) and electrooculography (EOG)). This increases familiarity with the procedure, laboratory and electrodes and is common use in sleep studies. Data was not recorded. Session three and four were the two experimental sessions which took place on the same weekday, spaced one week apart. The sessions started around 9 p.m. with some questionnaires and the attachment of the electrodes. Before sleep, we confronted subjects with three memory tests (semantic verbal fluency, SVF; Regensburger Wortflüssigkeitstest, RWT, and a paired-associate learning task, PAL) which were repeated in the morning. Afterwards, subjects were allowed to go to bed and were instructed to listen to the tape that was played after closing the door of the experimental room. Although subjects were asked to listen to the tape they were allowed to fall asleep whenever they wanted. Either the hypnosis or the control tape was played in sessions three and four in a randomized order. Eight hours after closing the door, the subjects were awakened. A vigilance test (psychomotor vigilance test, PVT) and a questionnaire on subjective sleep quality (Schlaffragebogen A, SF-A [24]) preceded the recall of the three memory tests. Afterwards, subjects were allowed to use the shower and were paid after the last session.

### Hypnotic and control tape

A German- and French-speaking hypnotherapist from the University of Fribourg (Laurent Rossier) translated the two texts we had previously used [14,15] to French. Duration of both texts was around 15 minutes. We played both texts with a comfortable volume via loudspeakers placed on a cupboard next to the bed. According to the German version, the hypnotic text was spoken slowly, with a soft, calm and gentle voice while the control text was read with a usual every-day voice and normal speed.

### Memory tasks

#### Word pair learning task (PAL)

We used the word pair learning task as described by Rasch et al. [25], while the words were translated to French. During learning, the word pairs were presented visually in black font on white ground on the screen, intermitted by a blank interval of 500 ms and a fixation cross (500 ms). Each word was displayed for 1000 ms. A blank screen of 200 ms separated the two words of a word pair. Subjects were asked to memorize as many words as possible. After 8 hours of sleep, subjects were awakened and recall was again tested by presenting the first word in black font on the white screen for 1000 ms, followed by a question mark after which subjects were asked to name the corresponding second word aloud or to say “next” in case they did not remember the word. The experimenter entered a 1 for correctly remembered words or a 0 for wrong or no answer. No feedback was given to the subjects. A blank screen preceded the following trial (500 ms). The order in which the word pairs were recalled was the same in both recall phases but differed from the order in which the pairs were learned.

Due to technical problems, presleep memory performance of 2 subjects and postsleep memory performance of one subject is missing. The performance level before sleep was on average 34.51 ± 1.95 (SEM) words out of 80 word pairs (43.14 ± 2.44%), indicating a medium task difficulty, excluding ceiling or floor effects. Memory performance after sleep was on average 33.95 ± 1.92 (SEM) words (i.e. 42.44 ± 2.49%). The relative performance level of postsleep recall was measured by setting presleep performance to 100%.

#### Semantic Verbal Fluency (SVF)

As in Cordi et al. [15], we applied the frontallobe-dependent longterm-memory task in which subjects have to name as many examples for a given category as possible within 2 minutes. Across the two experimental sessions and the immediate and postsleep recall, we randomized four categories across subjects (hobbies, fruits, animals, occupation). Double namings as well as words with the same word stem were excluded. Memory performance was measured by counting all valid examples that were written down. Relative improvement was calculated by setting presleep performance to 100%.

#### Regensburger Wortflüssigkeitstest (RWT)

Similarly, in the letter-based version of this task, subjects are given a letter and have to come up with as many words that start with this letter as possible within 2 minutes. Again, we randomized four letters across the four measure times across subjects (letters T, N, I, R). We set presleep performance to 100% and analyzed changes across sleep in reference.

### Questionnaires

#### Harvard Group Scale of Hypnotic Suggestibility (HGSHS)

In this standardized test to measure the hypnotizability of subjects, a hypnotic text is played from a tape leading subjects into a state of trance in which motoric, perceptual, cognitive and posthypnotic suggestions are given. Afterwards, subjects fill out a questionnaire asking for their subjective experiences during listening. According to the depth of reported hypnotic state during the whole procedure, subjects were divided into a high and low suggestible group (cutoff value 7). The high suggestible group (9 males, 10 females) on average achieved a score of 7.36 ± 0.16, while low suggestibles scored 4.65 ± 0.26 (10 males, 14 females). Those means differed significantly (*t*(41) = 8.26, *p* < .001). The two groups did neither differ in age (*p* > .30) nor gender distribution (*p* > .70). The German version of this test was translated into French and recorded by Laurent Rossier.

#### Schlaffragebogen Version A (SF-A)[24]

To measure subjective sleep quality, subjects filled out the SF-A in the morning. Sleep quality was assessed by averaging the ratings in question 13 in which participants had to judge the quality of the night-time sleep on 5 adjectives. Scorings were coded in a way that higher values indicate better sleep quality. Data of one subject is missing in this analysis.

#### Pittsburgh Sleep Quality Index (PSQI) [26]

With this questionnaire, subjects indicated for the last four weeks before participation how they rate their general sleep quality.

### Polysomnographic recordings

Electromyographic (EMG), electrocardiographic (ECG), electrooculographic (EOG), and electroencephalographic (EEG) electrodes were attached for polysomnography. Impedances did not exceed 10 kΩ. EEG was recorded with a 32-channels Easycap Net (Easycap GmbH, Herrsching) and a sampling rate of 500 Hz. At preprocessing with Brain Vision Analyzer 2.1 (Brain Products, Gilching, Germany), the electrodes were referenced against the mean of the mastoids (electrodes Tp9 and Tp10) for sleep data and against the mean of all electrodes for analyzing EEG activity during listening to the tapes. For scoring, data was filtered according to the settings suggested by the American Association of Sleep Manual (AASM)[27]. Two independent sleep scorers visually scored 30 second segments of sleep to define the stages 1-3, REM sleep, and wakefulness offline and according to standard criteria [27]. Derivations F4, C4, O2, HEOG, VEOG, and EMG were used therefore. A third sleep expert was consulted in case of disagreement. All scorers were blind to condition.

### EEG data analysis

EEG was preprocessed using Brain Vision Analyzer 2.1 (Brain Products, Gilching, Germany). Data was filtered (low-pass filter: 0.1 Hz; high pass filter: 50 Hz, Notch filter: 50 Hz). For spectral analysis of NREM sleep, only segments scored as S2 and S3 were selected. Artefacts were excluded. Before applying the Fast Fourier Transformation with a Hamming window of 10%, we further segmented data into ∼4 second segments with 10% overlap (4096 data points, 409 overlap). We exported area information on the power values of slow-wave activity (SWA, 0.5 - 4.5 Hz), theta (4.5 - 8 Hz), alpha (8 - 11 Hz), slow (11 - 13 Hz) and fast spindles (13 - 15 Hz). We defined 6 regions (frontal, central, parietal electrodes, left and right side respectively) to test electrode assemblies on power differences between condition and group. To control for possible total power differences between the two experimental sessions within the subjects, we calculated the relative power of the respective frequency band with total Power (0.5 - 50 Hz) set to 100%. Thus, the analyzed values are percentages of total power.

### Statistical analysis

The data was analyzed using paired-samples t-tests or analyses of variance with the between-subjects factor group (high vs. low) and the within-subjects factor text (hypnosis vs. control). In additional analyses we also included the within-subjects factors hemisphere (left vs. right) and topography (frontal vs. central vs. parietal). Follow-up t-tests were performed with paired-samples t-tests within the groups or with t-tests for independent samples. The level of significance was set to *p* = .05. Degrees of freedom were adjusted when homogeneity of variances was not given.

## Results

### Sleep stages

As expected, high suggestible subjects spent significantly more minutes in SWS after hypnosis (137.34 ± 9.74 min) than after the control text (124.21 ± 9.03 min; *t*(18) = 2.22, *p* = .04, effect size d = 0.32), see Fig 1B. The absolute difference was ca. 13 minutes, which reflects a 10.5% increase in night-time SWS. In low suggestibles, this difference was non-significant (124.29 ± 8.27 min vs. 134.21 ± 8.85 min, *p* = .11). This interaction between group and text was significant in a repeated measures ANOVA with *F*(1, 41) = 7.32, *p* = .01, eta^2^ = 0.15. It was not significant for any other sleep stage (all *p* > .30, see Table 1).

**Table 1.** Effects of group and text on the sleep stages

| Sleep stage | High suggestibles (n = 19) |  | Low suggestibles (n = 24) |  | *p*<br>inter-<br>action |
| --- | --- | --- | --- | --- | --- |
|  | Hypnosis | Control | Hypnosis | Control |  |
| Wake (min) | 9.66 ± 2.95 | 7.47 ± 2.35 | 15.29 ± 4.08 | 13.79 ± 4.36 | .93 |
| N1 (min) | 17.42 ± 2.38 | 16.24 ± 1.91 | 19.42 ± 2.22 | 18.04 ± 1.68 | .96 |
| N2 (min) | 213.03 ± 7.51 | 225.50 ± 7.84 | 198.40 ± 8.95 | 199.94 ± 7.85 | .34 |
| N3 (min) | **137.34 ± 9.74\*** | **124.21 ± 9.03** | 124.29 ± 8.27 | 134.21 ± 8.85 | **.01\*** |
| % N3 min increase<br>(control set to 100%) | 110.51 ± 34.19 | 100 ± 31.70 | 92.61 ± 30.18 | 100 ± 32.31 | **.01\*** |
| REM (min) | 86.95 ± 6.48 | 84.82 ± 5.91 | 83.96 ± 4.12 | 78.98 ± 5.69 | .75 |
| % Wake | 2.07 ± 0.64 | 1.64 ± 0.52 | 3.71 ± 1.06 | 3.08 ± 0.99 | .91 |
| % N1 | 3.74 ± 0.52 | 3.55 ± 0.42 | 4.49 ± 0.56 | 4.08 ± 0.39 | .79 |
| % N2 | **45.79 ± 1.62\*** | **49.13 ± 1.73** | 44.41 ± 1.65 | 44.77 ± 1.69 | .24 |
| % N3 | 29.47 ± 2.08<sup>+</sup> | 27.02 ± 1.93 | 27.77 ± 1.67 | 29.81 ± 1.80 | **.015\*** |
| % N3 increase<br>(control set to 100%) | 109.06 ± 7.68<sup>+</sup> | 100 ± 7.14 | 93.14 ± 5.59 | 100 ± 6.04 | **.014\*** |
| % REM | 18.66 ± 1.37 | 18.35 ± 1.20 | 18.85 ± 0.85 | 17.71 ± 1.28 | .68 |
| Total sleep time (min) | 465.53 ± 3.31 | 459.55 ± 4.57 | 444.69 ± 7.04 | 447.33 ± 6.29 | .30 |
| Sleep latency (min) | 15.21 ± 2.84 | 18.71 ± 4.11 | 33.79 ± 6.50 | 29.77 ± 5.00 | .34 |
| SWS latency (min) | 15.32 ± 1.73 | 16.03 ± 1.60 | **12.17 ± 0.95\*** | **15.48 ± 1.66** | .30 |
| REM latency (min) | 137.79 ± 12.43 | 111.84 ± 8.90 | 111.33 ± 9.10 | 124.56 ± 11.09 | .06<sup>+</sup> |
| Sleep efficiency | 0.97 ± 0.01 | 0.96 ± 0.01 | 0.93 ± 0.02 | 0.94 ± 0.01 | .30 |
| Subjective sleep quality | 3.77 ± 0.14 | 3.53 ± 0.17 | 3.41 ± 0.19 | 3.68 ± 0.17 | **.05\*** |

Displays means in % and min ± SEM. The right column indicates *p* values for the group\* text interaction. Bold values indicate within group effects. Asterisks indicate significant differences with \*: *p* < .05, <sup>+</sup>: *p* ≤ .09. Sleep efficiency: number of minutes of sleep divided by number of minutes in bed.

**Figure 1.**
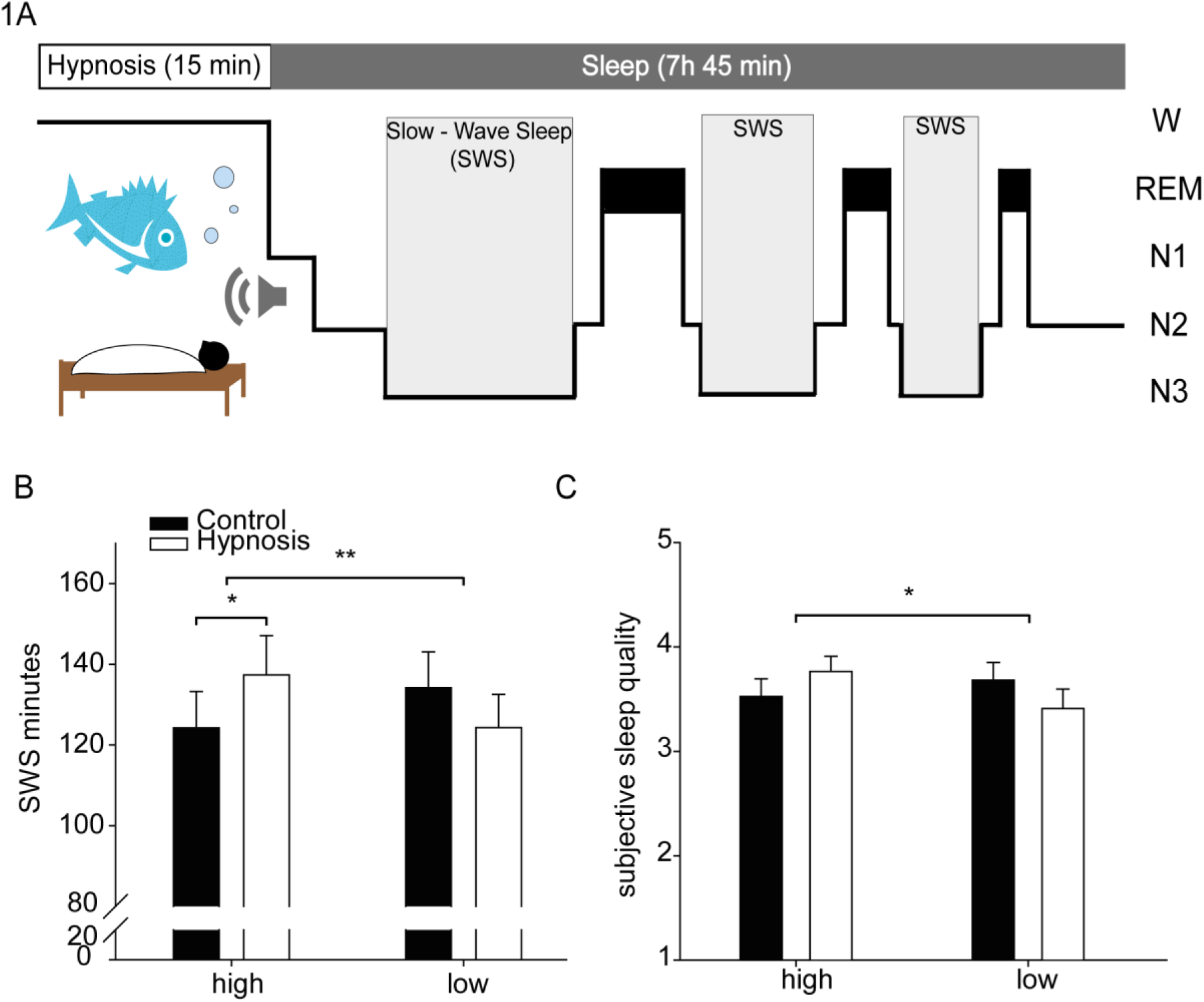
Session flow and main results. Session flow and main results. (A) Healthy young participants listened either to a hypnosis or a control tape in the evening before sleep in a cross-over design. They were allowed to sleep for a full night of sleep afterwards. Sleep was measured using polysomnography including 32 EEG channels. (B) High suggestible participants spent significantly more time in slow-wave sleep (SWS) as compared to the control text. No effect was observed in low suggestibles. The interaction was significant (*p* < .01). C) Subjective evaluations of sleep quality followed the similar pattern as the time spent in SWS (Interaction *p* < .05). Means +/- standard errors of the means (SEM) are indicated. *: *p* < .05. **: *p* ≤ .01.

Also for the percentages of the sleep stages, repeated measure ANOVAs confirmed a significant group * text interaction only for percentage of slow-wave sleep (*F*(1, 41) = 6.51, *p* = .015, eta^2^ = 0.14, all other interactions *p* ≥ .06, see Table 1). High suggestibles tended to have more % SWS after the hypnosis tape (29.47 ± 2.08%) than after the control tape (27.02 ± 1.93%), *t*(18) = 1.85, *p* = .08 (see Table 1), while descriptively the opposite was true for low suggestibles (29.81 ± 1.80% after control tape, 27.77 ± 1.67% after hypnosis, *t*(23) = -1.75, *p* = .09).

Interestingly, we also observed a statistical trend for an influence of hypnosis on REM sleep latency (*F*(1, 41) = 3.69, *p* = .06, eta^2^ = 0.08). In high suggestibles, REM latency was descriptively longer after hypnosis as compared to the control condition, while the opposite pattern occurred for low suggestibles (see Table 1). All other interactions with objective sleep parameters were neither significant nor a trend (*p* > .10). In addition, a main effect of group was observed for sleep latency (*F*(1, 41) = 5.87, *p* = .02, eta^2^ = 0.13 and sleep efficiency (*F*(1, 41) = 5.26, *p* = .03, eta^2^ = 0.11), (all others *p* > .10). High suggestibles fell asleep earlier (16.96 ± 2.79 min) than low suggestibles (32.03 ± 5.01 min), *t*(35.15) = 2.63, *p* = .013, effect size d = 0.75 and had a higher ratio of sleep efficiency (0.96 ± 0.01) than low suggestibles (0.93 ± 0.01), *t*(34.89) = -2.47, *p* = .02, d = 0.60. No main effect of text was significant (all *p* > .10).

### Process across the first 6 hours of sleep

In order to display the time course of hypnotic influence on sleep across the night, we extracted the first six hours of sleep for each person from sleep onset onwards, defined as first N1 which was followed by N2 (see Fig 2A). The ANOVA with group as between-subjects factor and time (hour 1 to 6) as within-subjects factor on the difference between hypnosis and control in SWS was non-significant (*p* > .30). The main effect of group was significant (*F*(1, 41) = 4.27, *p* = .045, eta^2^ = 0.09) with a larger difference in high suggestibles (1.54 ± 1.14 min) than low suggestibles (-1.43 ± 0.91 min), *t*(41) = -2.07, *p* = .045, presumably reflecting the interaction found for the amount of overall SWS.

**Figure 2.**
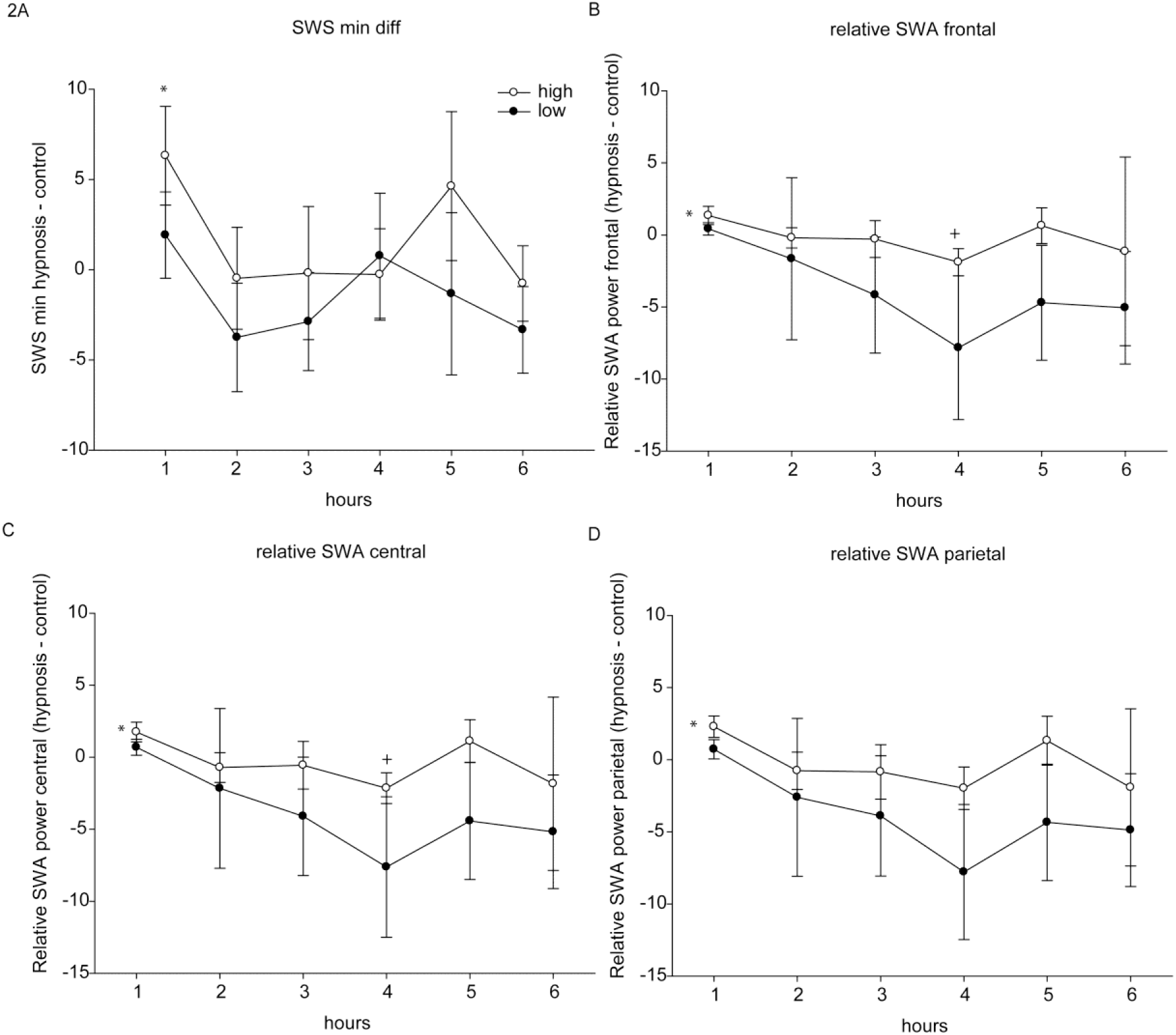
Process of minutes SWS and SWA power across the first 6 hours of sleep. Fig 2. displays the process across the first 6 hours of sleep in high (white dots) and low suggestible (black dots) subjects. (A) SWS minutes in hypnosis minus control. The difference in SWS between the groups did not differ at any of the single time points (*p* > .20). In highly suggestible subjects the value in the first hour was however significantly different from zero (*t*(18) = 2.31, *p* = .03). No other values differed from zero in this (all *p* > .20) or the other group (all *p* > .10). *(B to D)* SWA power after hypnosis minus control in frontal, central and parietal electrodes. The group difference was not significant at any of the single time points (*p* ≥ .10). In highly suggestible subjects the overall value in the first hour was however significantly different from zero (*t*(18) = 2.64, *p* = .02). No other values differed from zero in this (all *p* > .07) or the other group (all *p* > .06).

Like for SWS we analyzed relative SWA power (0.5 – 4.5 Hz) in NREM sleep across the first six hours of sleep after sleep onset to investigate sleep depth development. In a repeated measures analysis in SWA across the six times points, the main effect of time was significant (*F*(3.7,150.5) = 10.80, *p* < .001, eta^2^ = 0.21). Here, all differences were highly significantly different from each other (all *p* ≤ .02) except from hours 2 and 3 (*p* = .60), 2 and 4 (*p* = .17), 2 and 5 (*p* = .10) and 4 and 5 (*p* = .99). The interaction between time and group was a trend (*F*(5, 205) = 2.01, *p* = .078, eta^2^ = 0.05). In follow-up analyses, groups did not significantly differ from each other in any of the hours (all *p* ≥ .18). All other main effects and interactions were non-significant (all *p* >.20).

We additionally analyzed the two groups separately across the 6 hours and found that the difference between hypnosis and control is significantly different from zero in high suggestibles in the first hour (*t*(18) = 2.64, *p* = .017). Further, a trend appeared in the fourth hour (*t*(18) = -1.82, *p* = .09). For low suggestibles all comparisons were *p* > .10. This pattern was found in frontal, central and parietal electrodes as reflected in Fig 2B-D.

In addition to SWA during NREM in the first six hours of sleep, we also calculated average oscillatory power of different frequency bands during NREM sleep of the whole night. In repeated measures ANOVAs we tested the within-subjects factors text, hemisphere, topography and between-subjects factor group on the relative power in several frequency bands. The main effect of text for relative SWA (*F*(1,41) = 4.52, *p* = .04, eta^2^ = 0.10) showed higher SWA after the hypnosis text (89.60 ± 0.44%) than after the control text (88.42 ± 0.55%) as confirmed in the follow-up t-test (*t*(42) = 2.04, *p* = .048). There was a group main effect (*F*(1,41) = 34.94, *p* = .03, eta^2^ = 0.11) with higher SWA in the low suggestible (89.78 ± 0.49%) than the high suggestible group (88.04 ± 0.62%), *t*(41) = 2.22, *p* = .03). Also, the three-way interaction between topography, text and group was significant (*F(*2,40) = 3.94, *p* = .03, eta^2^ = 0.16). Follow-up t-tests in high suggestibles revealed a trend in the effect of text (*t*(18) = 2.04, *p* = .057), with higher SWA after hypnosis (88.92 ± 0.64%) than after control (87.16 ± 0.86%), a main effect of topography (*F*(2,17) = 51.69, *p* > .001, eta^2^ = 0.86) and a significant interaction between both factors (*F*(2,17) = 7.95, *p* = .004, eta^2^ = 0.48). SWA after hypnosis was significantly higher than after control for frontal and parietal electrodes (*t*(18) = 2.22, *p* = .039, *t*(18) = 1.55, *p* = .14, *t*(18) = 2.18, *p* = .043 for frontal, central and parietal electrodes, respectively). In low suggestibles, only a main effect of topography was significant (*F*(2,22) = 134.61, *p* >.001, eta^2^ = 0.92). Also when analyzing the relative increase with SWA power after control set to 100% the main effect of text was significant (*F*(1,41) = 4.59, *p* = .04, eta^2^ = 0.10) with higher SWA after hypnosis (101.33 ± 0.50%) than after control (100 ± 0.59%), *t*(42) = 2.05, *p* = .047.

For relative theta power (4.5 – 8 Hz), no main effects or interactions with group or text appeared (*p* > .07). We neither found main effects or interactions with text for slow spindles (11 - 13 Hz; *p* > .20) nor fast spindles (13 - 15 Hz; *p* > .20). There was a main effect of group for relative alpha power (8 – 11 Hz) (*F*(1,41) = 4.58, *p* = .04, eta^2^ = 0.10) with higher alpha power in high (2.51 ± 0.20%) than low suggestibles (2.02 ± 0.13%), *t*(41) = -2.14, *p* = .04.

The amount of SWA did not correlate with the amount of SWS (*p* > .10). Neither did the difference between SWA after hypnosis vs control correlate with the difference in SWS between the two conditions (*p* > .20).

### Power analyses during the suggestion

We performed the same analyses as during NREM sleep for the time in which the subjects listened to the suggestion before falling asleep. In this time frame, relative theta activity was higher during hypnosis (13.33 ± 0.58%) than during the control text (11.47 ± 0.55%), *F*(1, 41) = 16.93, *p* = < .001, eta^2^ = 0.29. The other interactions with group or text were *p* ≥ .19. The four-way interaction between topography, hemisphere, text and group was a trend (*F*(2, 82) = 2.93, *p* = .06, eta^2^ = .07). Only in parietal electrodes the three-way interaction hemisphere, text and group was significant (*F*(1, 41) = 5.01, *p* = .03, eta^2^ = 0.11). Text* group interactions were however neither significant in only right or left parietal electrodes (*p* > .20) neither were group differences (*p* > .10).

No main effects for text or griup were significant for relative SWA (*p* > .50) or alpha (*p* ≥ .20) during listening. For SWA a significant interaction between topography, text and group appeared (*F*(2,40) = 4.30, *p* = .02, eta^2^ = 0.18). In the follow-up tests, there was only one significant trend in the interaction between topography and text (*F*(2,17) = 3.34, *p* = .06, eta^2^ = .28), but further tests did not reveal significant differences between hypnosis and control condition for any of the regions (all *p* > .60)

Theta power during listening correlated negatively with later SWA power across the whole night (*r*(42) = -.32, *p* = .035). No outliers were detected here. The power difference in theta between hypnosis and control condition did neither correlate with later difference in SWA (*p* >.10 across the whole night) nor with the difference in the amount of SWS (*p* > .60). These correlations were also true excluding one extreme outlier as defined by more than three lengths of the height of the according boxplot.

### Subjective sleep quality and covariates

Subjective sleep quality ratings differed depending on group and text (*F*(1,40) = 4.09, *p* = .05, eta^2^ = 0.09). Although post-hoc t-tests were neither significant in high suggestibles (hypnosis 3.77 ± 0.14, control 3.53 ± 0.17, *t*(18) = 1.21, *p* = .24) nor low suggestibles (hypnosis 3.41 ± 0.19, control 3.68 ± 0.17, *t*(22) = -1.69, *p* = .11), descriptive data follows the same direction as the objective measures of SWS. Thus, hypnosis rather improved subjective sleep quality in high, but not low suggestible subjects. We did not find a correlation between the difference in subjective ratings between hypnosis and control and the difference in the amount of SWS between both conditions (*p* > .80).

Vigilance as measured by PVT reaction times did not differ between the two nights (*p* >.60). As Table 2 shows, neither gender nor sleep quality as defined by the PSQI value of below 5 versus 5 and higher differed between those people who benefited from hypnosis for SWS and those who did not.

**Table 2.** Relationship between positive effect of hypnotic suggestions and covariates.

|  | SWS after hypnosis higher | SWS after control text higher | Chi<sup>2</sup> p |
| --- | --- | --- | --- |
| Total | 25 | 18 |  |
| Male/female | 12/13 | 7/11 | .55 |
| PSQI <5/≥ 5 | 11/14 | 10/8 | .46 |

Displays the distribution of subjects that showed increased or decreased SWS amount after hypnosis across several dimensions.

## Memory performance

*PAL*. Percentage of improvement across the night was significantly influenced by the text preceding the sleep period (*F*(1,38) = 6.36, *p* = .02, eta^2^ = 0.14). Performance across a night of sleep after hypnosis decreased (94.61 ± 2.03%) compared to the night after the control text (100.96 ± 2.23%), *t*(39) = -2.64, *p* = .01. This effect was however mainly driven by low suggestible subjects (*p* = .02, versus *p* = .30 in high suggestibles).

*RWT*. Memory performance before sleep was on average 16.77 ± 0.77 words, after sleep 16.22 ± 0.82 words. No main effects or interactions of text or group appeared (all *p* > .50).

*SVF*. Memory performance before sleep was on average 21.37 ± 0.86 words, after sleep 19.98 ± 0.79 words. No main effects or interactions of text or group appeared (all *p* > .20), see Table 3.

**Table 3.** Memory performance in high and low suggestible subjects in both text conditions

|  | high suggestibles<br>(PAL: n = 18, others: n = 19) |  | low suggestibles<br>(PAL: n = 22; others: n = 24) |  | *p*<br>inter-<br>ac-<br>tion |
| --- | --- | --- | --- | --- | --- |
|  | hypnosis | control | hypnosis | control |  |
| PAL abs. presleep | 32.78 ± 2.99 | 34.50 ± 3.73 | 35.23 ± 1.99 | 32.82 ± 2.60 | .07 |
| PAL abs. postsleep | 31.50 ± 3.10 | 35.22 ± 3.98 | 33.68 ± 2.36 | 33.64 ± 2.59 | .16 |
| PAL % presleep | 40.97 ± 3.73 | 43.13 ± 4.66 | 44.03 ± 2.50 | 41.02 ± 3.25 | .07 |
| PAL % postsleep | 39.38 ± 3.87 | 44.03 ± 4.98 | 42.10 ± 2.95 | 42.05 ± 3.24 | .16 |
| PAL improvement (presleep = 100%) | 94.96 ± 2.38 | 98.65 ± 4.24 | **94.33 ± 3.18\*** | **102.85 ± 2.12** | .32 |
| RWT improvement (presleep = 100%) | 112.26 ± 13.32 | 108.35 ± 19.77 | 114.96 ± 10.68 | 104.17 ± 7.67 | .76 |
| RWT | -.053 ± 1.92 | -2.00 ± 1.97 | 0.08 ± 1.49 | -0.38 ± 1.25 | .61 |
| SVF improvement (presleep = 100%) | 93.28 ± 6.92 | 105.94 ± 11.42 | 100.17 ± 5.90 | 106.00 ± 8.96 | .65 |
| SVF | -1.84 ± 1.65 | -1.11 ± 2.59 | -0.92 ± 1.11 | -1.00 ± 1.69 | .79 |

Displays absolute and relative values for the three memory tests that were performed, separately for high and low suggestible subjects for both text conditions. The right column indicates *p* values for the group \* text interaction. Asterisks indicate significant differences \**p* < .05.

### Correlations between memory and sleep

Only in low suggestibles we found a significant correlation between the difference in the amount of SWS and the difference in the RWT *r*(23) = .46, *p* = .02 (*r*(23) = .42, *p* = .04 for difference in SWS minutes and RWT). Also a trend for a correlation between the difference in percent SWS and an improvement in PAL resulted in this sample (*r*(21) = .39, *p* = .07). With SVF improvement, this correlation was not significant (*r*(23) = .07, *p* = .76. In high suggestibles, the correlations between the difference in SWS and PAL was *r*(17) = -.21, *p* = .40, with RWT *r*(23) = .04, *p* = .88 and with SVF *r*(23) = -.26, *p* = .29.

## Discussion

Slow-wave sleep is an important component of healthy sleep and plays a vital role for mental health. Here we show that listening to hypnotic suggestions before sleep significantly increased the amount of SWS in highly suggestible young subjects even across a whole night of sleep. The duration of objectively measured SWS increased by 13 minutes in highly suggestible subjects, and these extensions in SWS also affected subjectively rated sleep quality. Our results are a conceptual replication of our two previous studies [14,15]. In these two studies, we reported that the same hypnotic suggestion increased SWS in a midday nap in young and old healthy and high-suggestible females. In our study in young participants, we had also excluded potential confounds of general relaxation properties of the hypnotic vs. the control text as well as demand characteristics [see control studies in, 14]. Interestingly, the hypnotic suggestion increased SWS amount during the nap on average by ca. 9 minutes, which is in a similar range as the 13 minutes reported here for night-time sleep. Thus, we successfully generalized the previously reported SWS benefit of hypnotic suggestions to night-time sleep, male participants and a different language (French).

Please note that our initial prediction was that hypnotic suggestions before a period of night-time sleep would have a lower effect on SWS duration as compared to our nap studies. As homeostatic sleep pressure is typically higher in the evening as compared to midday for young and healthy non-habitual nappers, we expected that effects of hypnotic suggestions on SWS duration might be smaller during night-time sleep. In contrast to our notion, the effects of hypnotic suggestions on SWS duration appear to be comparable between naps and night-time sleep, at least in absolute terms. Due to the high differences in total amount of SWS in naps and night-time sleep, results differ strongly in relative terms (i.e., 50-80% increase of SWS by hypnotic suggestions in naps vs. 9% in night-time sleep).

As expected from the nap studies, we observed the strongest effects of hypnotic suggestions on SWS duration in the first hour of night-time sleep in highly suggestible participants. On the descriptive level, some hypnosis-induced increases in SWS were also observed in the fifth hour, but this difference did not reach significance. Also for oscillatory activity in the slow-wave range (SWA), effects of hypnotic suggestions were most visible in the first and possibly the second hour of night-time sleep as well as in the fourth hour. The increase of SWA by hypnosis was broadly spread and affected frontal, central and parietal recording sites.

Also similar to our two previous reports, we found no or even opposite effects of hypnotic suggestions on SWS and SWA in low suggestible participants. As discussed previously [15], opposite result patterns for low as compared to high suggestible participants have been observed also for other type of hypnotic suggestions [see e.g., 28,29]. The reasons are not entirely clear, but it might be possible that at least some low-suggestible participants actively counteract the intended direction of the hypnotic suggestions, possibly due to a fear of being hypnotized and/or losing self-control [29]. Interestingly, the strongest decrease in SWS duration and SWA after the hypnotic suggestions occurred not in the first, but in second hour of sleep. One could speculate that the homeostatic sleep pressure in the first hour of night-time sleep is so dominant that a counteracting of hypnotic suggestions with the aim to reduce SWS is only successful after some SWS need is fulfilled. However, future studies are needed to examine this speculation.

We also analyzed the power in the frequency bands during listening to the hypnotic suggestions and compared them to the activity during listening to the control text. We observed that both, high and low suggestible subjects showed increases in theta during the hypnotic compared to the control text. Some previous studies have as well reported theta band specific increases accompanying the hypnotic state [30]. Most of them have however stated that this increase should be higher for highly suggestible subjects. Moreover, highly suggestible subjects have previously shown more theta activity not only during hypnosis, but also during control conditions [30,31] which was neither the case in our data.

Unexpectedly, the observed increase in theta power during listening was negatively correlated to the amount of SWA observed during the NREM periods of sleep, which also contrasts previous reports [e.g., 22]. Furthermore, increases in theta oscillation during listening were not related to the effect of hypnotic suggestions on later SWS, as reported in our previous nap studies. One problem could be that theta oscillations are not a specific marker for the hypnotic state, but are also related to general mental relaxation procedures as well as tiredness and sleepiness. This might be particularly problematic in times of increase sleep pressure (i.e., in the evening before night-time sleep).

Contrary to other studies showing that increasing SWA or SWS positively influences memory performance, we did not observe any changes in memory across deepened sleep. We did neither find a benefit in the episodic nor the semantic memory task. It has been discussed before that possibly, a certain presleep performance level or memory strength is needed for a beneficial effect of sleep [32,33]. Possibly, the performance level in our sample of only about 40% of the presented word pairs was too low to achieve a sleep-dependent benefit. For the verbal fluency task, we also did not observe any positive effect of extending SWS by hypnotic suggestions as in our nap study in younger adults. A positive effect in this task appears to be specific to older adults.

Limitations. It could be argued that instead of increasing sleep depth, the control tape has reduced sleep quality compared to a normal night of sleep. As we did not measure sleep without presenting any tape, we have no neutral baseline to fully exclude this alternative explanation. Thus, we cannot prove whether the control tape had influenced sleep patterns. Our method ensures that we do not include any unspecific effects of listening into the effect of hypnotic suggestions. It would however be interesting to include a third intervention-free night to exclude that listening to the control tape reduces sleep quality.

In sum, hypnotic suggestions are effective to increase the amount of deep sleep in a young, healthy sample of good sleepers even across a normal night of sleep. This demonstrates the massive effect of hypnotic suggestions as a highly functional, healthy sleep pattern could still be influenced. Also on the subjective level, subjects’ rating of their sleep quality followed this pattern. For practical reasons the effect should now be tested for patients with sleep problems. In addition, it is now very important to carefully examine and advance our theoretical understanding of the potential mechanism underlying the positive effect hypnotic suggestions on slow-wave sleep, which would stimulate further insight into this method and how it can be further improved.

## Acknowledgements

We thank Kamila Giruc, Corentin Wicht, Viviana Leupin, Nathalie Bürdel and Milica Vasic for assistance in data collection.

## Notes

The authors declare that they have no conflict of interest

